# Evotuning protocols for Transformer-based variant effect prediction on multi-domain proteins

**DOI:** 10.1101/2021.03.05.434175

**Authors:** Hideki Yamaguchi, Yutaka Saito

## Abstract

Accurate variant effect prediction has broad impacts on protein engineering. Recent machine learning approaches toward this end are based on representation learning, by which feature vectors are learned and generated from unlabeled sequences. However, it is unclear how to effectively learn evolutionary properties of an engineering target protein from homologous sequences, taking into account the protein’s sequence-level structure called domain architecture (DA). Additionally, no optimal protocols are established for incorporating such properties into Transformer, the neural network well-known to perform the best in natural language processing research. This article proposes DA-aware evolutionary fine-tuning, or “evotuning”, protocols for Transformer-based variant effect prediction, considering various combinations of homology search, fine-tuning, and sequence vectorization strategies. We exhaustively evaluated our protocols on diverse proteins with different functions and DAs. The results indicated that our protocols achieved significantly better performances than previous DA-unaware ones. The visualizations of attention maps suggested that the structural information was incorporated by evotuning without direct supervision, possibly leading to better prediction accuracy.

**Availability:** https://github.com/dlnp2/evotuning_protocols_for_transformers

**Supplementary information:** Supplementary data are available at *Briefings in Bioinformatics* online.

## Introduction

A number of mutagenized proteins obtained by evolutionary engineering methods [1] have been reported to demonstrate highly improved functional activity in biomedical research: to name a few, fluorescence [2], signal transduction [3], or base editing capability [4]. It is well-known that many natural proteins have multiple domains, whose functions are determined by the serial arrangement of domains called domain architecture. For example, approximately 65% of eukaryotic proteins are considered to be multi-domain [5]. Furthermore, various industrially essential proteins, such as artificial antibodies [6] or CRISPR/Cas system-related proteins [7], have multiple domains. In protein engineering, a specific domain in a multi-domain protein is often mutagenized (e.g., [8-17]).

While useful, the mutagenesis experiments are often costly because multiple iterations of library construction and selection are necessary and they usually require target-specific human knowledge, the latter of which restricts the methods’ generalizability to other proteins. In recent years, machine learning techniques are utilized to predict variant effects for tackling the challenges [18-20]. Among them, more attentions are paid to an approach called representation learning, in which features (or descriptors) are directly learned from primary sequences alone [21-23], inspired by natural language processing (NLP) research. The approach is promising because 1) it can exploit tens of millions to billions of sequences in public databases without the necessity for mutagenesis experiments or human expertise; 2) it automatically generates a “contextualized” feature for each amino acid in a given sequence explicitly considering interactions between residues in contrast to “static” features that take fixed values depending only on the type of amino acids [24-26].

In this direction, a representation learning protocol named “evotuning” (evolutionary fine-tuning) was proposed to incorporate evolutionary properties of proteins into LSTM [27] based representation learning models, whose performances were evaluated through variant effect prediction tasks [28]. In the protocol, the models were first trained using a large set of proteins across various families (pre-training). Next, homology search was performed with the full-length sequences of an engineering target protein as the queries to collect homologs. Then, the hit sequences were used for fine-tuning. Lastly, the full-length protein sequences were fed into the fine-tuned models to be vectorized (embedding). The evotuned models were demonstrated to perform significantly better than the pre-trained ones or models with static features.

This protocol, however, had been evaluated only on single-domain proteins. Therefore, it is still unclear whether the protocol is also successfully applied to proteins with multiple domains. For instance, let us consider that a full-length sequence of a multi-domain protein is used as a homology search query as conducted for a single-domain protein. This will result in sequences with partial homology to the query (i.e., aligned with some, but not all, of the multiple domains in the query). It is not obvious whether we should incorporate these sequences into fine-tuning to improve variant effect prediction performance. Thus, optimal protocols for multi-domain proteins need to be devised, particularly considering the issue of partial homology. Furthermore, to the best of our knowledge, none of the previous studies, including the aforementioned one, examined how such protocols affect the learned representations by Transformer [29], which is the neural network architecture known to achieve state-of-the-art performances in most NLP tasks [30, 31]. Also, Transformers explicitly calculate dependencies between residues known as attention map, which gives us clues to interpret how a prediction result for a specific variant is obtained.

Here we propose effective evotuning protocols for Transformer-based variant effect predictors, in which we explicitly consider domain architectures by combining homology search, fine-tuning, and embedding strategies. For evaluation, we systematically compared the models’ performances trained with our protocols on diverse proteins with different functions and domain architectures. The results demonstrated that our protocols achieved significantly better prediction accuracy compared to the previous one. Moreover, the visualization of attention maps in Transformer suggested that evotuning can incorporate structural information of the proteins without direct supervision, possibly leading to improved performances.

## Materials and methods

### Experimental pipeline

We outline the 4-stage experimental pipeline used in this study (Figure 1): (a) pre-training. A Transformer model was trained from scratch with about 32 million primary sequences in Pfam [32] to capture general properties of proteins without reference to the domain annotation data; (b) fine-tuning the pre-trained model with homologous sequences for each target protein to incorporate evolutionary information specific to the target; (c) embedding each wild-type or mutagenized sequence through the fine-tuned model to obtain the vectorized feature representation of the whole sequence. These vectors were used in the next step; (d) training and evaluating variant effect predictors with the measured variant effects to model sequence-function relationships regarding the features as the surrogates of protein sequences. Note that a similar experimental pipeline was also used in the previous study to evaluate the performance of an LSTM-based representation learning model and its evotuning for single-domain proteins [28].

**Figure 1.**
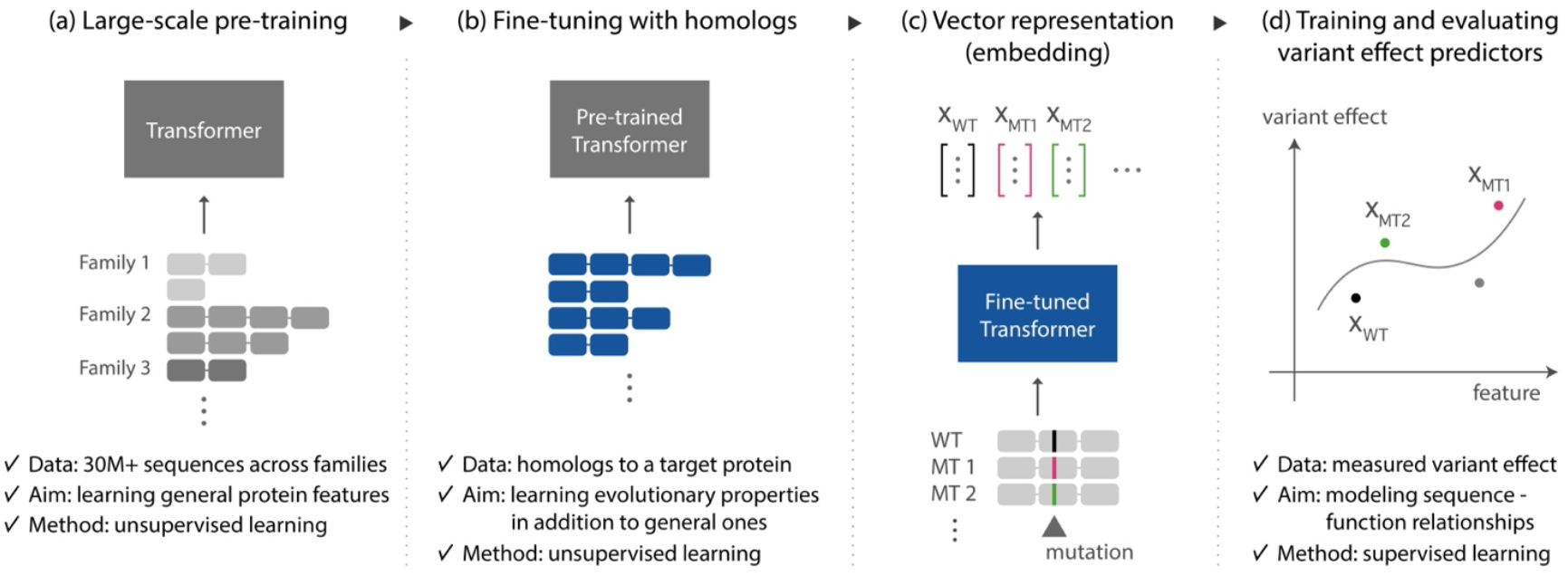
Overview of our 4-stage pipeline for protein representation model training and evaluation.

Because our primary interest was in elucidating how to utilize evolutionary information specific to a mutagenesis target protein effectively, the same pre-trained model was used in (a) for all experiments.

### Neural network implementation and training

We used a PyTorch [33] implementation of Transformer and a pre-trained weight reported in [21]. The pre-trained model’s perplexity, which measures its generalization, was 13.04 on Pfam holdout dataset [21]. The network architecture is the same as the encoder part with N=12 described in the Figure 1 in [29]: 12 multi-head attention layers with 12 heads for each. Each fine-tuning was performed on 4 NVIDIA V100 GPUs using Adam optimizer [34] with learning rate 0.0001, automatic mixed precision [35] enabled, and appropriate gradient accumulation steps for the whole batch to fit into the GPU memory.

We performed each training with masked language modeling (LM) [36] objective. In masked LM, some randomly selected amino acids in an original protein sequence are replaced by “<mask>” token before input to a Transformer model. Then, the model is trained to predict the original amino acids on the corrupted positions given information from all other residues. The loss is a cross entropy between the predicted and the original amino acids (Supplementary Figure S1). In this study, the token corruption probabilities were set to be the same as in [36]. The special tokens ([CLS] and [SEP]) used in masked LM were appended to the first and the last positions of each input sequence, respectively. Each sequence was then batched before fed into the network with batch size 256.

A learnable lookup table vectorized every amino acid before input into the neural network. Each training process was terminated when no improvement in validation loss was seen after ten epochs. We split the input sequences into training and validation sets by randomly selecting 90% and 10% sequences. Note that this splitting of training and validation sets was applied to homologous sequences used in fine-tuning (Figure 1b); the splitting in the evaluation of variant effect predictors (Figure 1d) was separately performed for variant sequences as described in the later section.

### Homology search

For fine-tuning, each homology search was performed with jackhmmer in HMMER software [37] against UniRef50 database [38]. The E-value threshold was set to 0.0001 for hit sequence inclusion (both --incE and --incdomE). Each run was stopped when converged or the iteration round reached 20. The sequences longer than 1.5x of the wild-type sequence length were removed. In addition to this length-based filtering, other strategies were also considered to remove noisy sequences, which will be described in a later section. We report the lists of domain architectures of the homologous proteins for each protein considered in this study (Table 1) in Supplementary Data S3. The majority of the homologous proteins had the same or similar domain architecture as in the query protein.

**Table 1.**
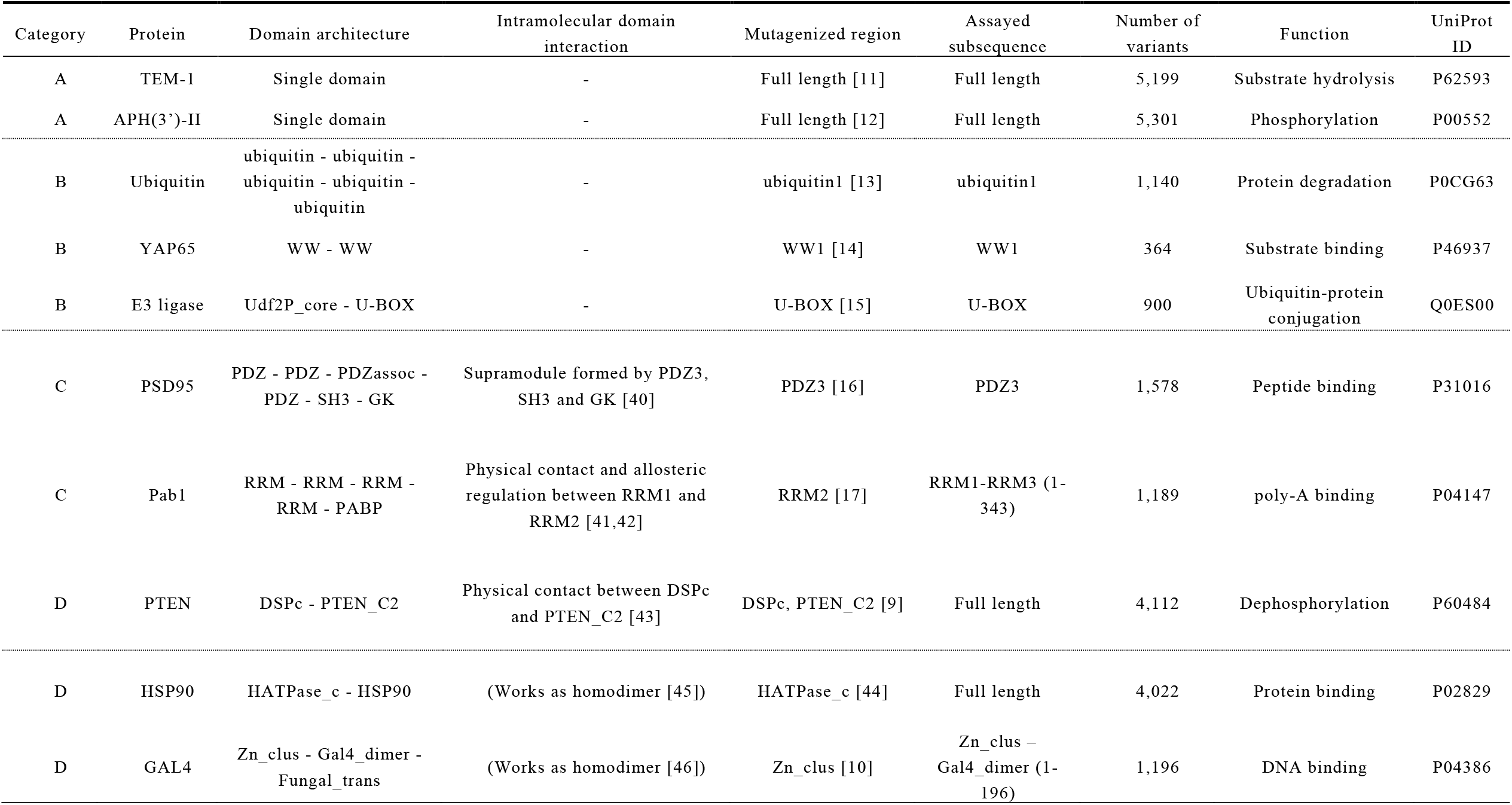
Summary of the examined proteins in this study. The domain architectures are based on Pfam [32]. In the “mutagenized region” column, the domain name followed by a number means its position in the domain architecture (e.g., PDZ3 in PSD95 means the third PDZ domain from the N-terminus). See the main text for the definition of the protein categories A-D.

### Domain annotation

For some of our protocols, we performed domain annotation on the hit sequences from homology search. We first run hmmscan in HMMER software using Pfam database as the reference. For each sequence, the outputs were parsed and filtered with the conditions [39]: 1) a domain candidate with an independent E-value larger than 0.0001 was removed; 2) if two domain candidates overlapped each other, the one with a smaller E-value was kept.

### Vector representation of protein sequences (embedding)

A sequence containing L amino acids was fed into a fine-tuned model to result in a matrix of the shape (L+2, 768), where 2 is for [CLS] and [SEP] tokens, and 768 is the embedding dimension. An average over the first dimension was taken to obtain a fixed-length vector representing the whole sequence, which was used as input to variant effect predictors in the later stage.

### Variant effect measurement data sourcing

We sourced the variant effect measurement data (i.e. protein sequence and measured value pairs) on ten different proteins [8-10] considering the following typical protein engineering scenarios: (A) the mutagenized protein has a single domain; (B) mutations are introduced to a specific domain in a multi-domain protein which is known to work independently on other domains; (C) similar to the scenario (B), but a specific domain is mutagenized which works cooperatively with other domains in a protein. In protein engineering, assay experiments to measure variant effects are often performed using a protein’s subsequence rather than the full-length sequence. Thus, we also collected the information on the assayed subsequences used in the original studies. Each mutation was a single amino acid substitution. The measured values were normalized according to the prescription described in [8].

In Table 1, the collected proteins, the mutagenized regions, and the assayed subsequences are summarized. Here we classified the proteins into four categories considering their domain architecture and (non-)existence of intramolecular domain interactions. The single-domain proteins (TEM-1 [11] and APH(3’)-II [12]) were classified into the category A; in the category B, the multi-domain proteins without intramolecular domain interactions were collected (ubiquitin [13], YAP65 [14], and E3 ligase [15]); the proteins known to have interacting domains fell into the category C (PSD95 [16, 40] and Pab1 [17,41,42]). Each of these protein categories corresponded to the respective protein engineering scenarios described above. We also prepared the category D for the proteins not considered to belong to the other categories: PTEN has two domains in physical contact with each other [43], but the mutations were introduced to both of the two interacting domains rather than one domain as in the category C [9]. HSP90 [44,45] and GAL4 [10,46] are known to work as homodimers, while the proteins in the category from A to C work as monomers. We here emphasize that the molecular functions of these proteins altered by the mutagenesis experiments are diverse: hydrolysis, substrate binding with distinct specificities, (de)phosphorylation, or functions related to protein degradation.

### Variant effect prediction

We used LASSO-LARS [47] implemented in scikit-learn Python package [48] as our variant effect predictor. LASSO-LARS is a sparse linear regression algorithm that is well-known for automatically selecting relevant features in high-dimensional space. In this study, the model was chosen to work well, particularly when the available number of measured variants is as small as a hundred or less. We note that the LASSO-LARS was also used in the previous study on evotuning for LSTM and single-domain proteins [28] for variant effect prediction. A separate predictor was trained for every combination of the selected protein and protocol. We examined two different training sizes (0.8 and 0.1) to emulate “high-N” and “low-N” scenarios, the latter of which is often the case in realistic protein engineering. We repeated the experiments using 32 different random seeds for splitting training/validation datasets to perform statistical tests for every combination of protein and protocol. The penalty coefficient on the L1 regularization term was determined by 10-fold nested cross-validation. The evaluation metric was Spearman’s correlation coefficient. The performance differences between protocols were tested with the one-sided Wilcoxon test using SciPy [49] Python package (scipy.stats.wilcoxon).

### Construction of the proposed protocols

In this work, we propose to construct the protocols as the combination of the options in the following three parts (Figure 2):

**Figure 2.**
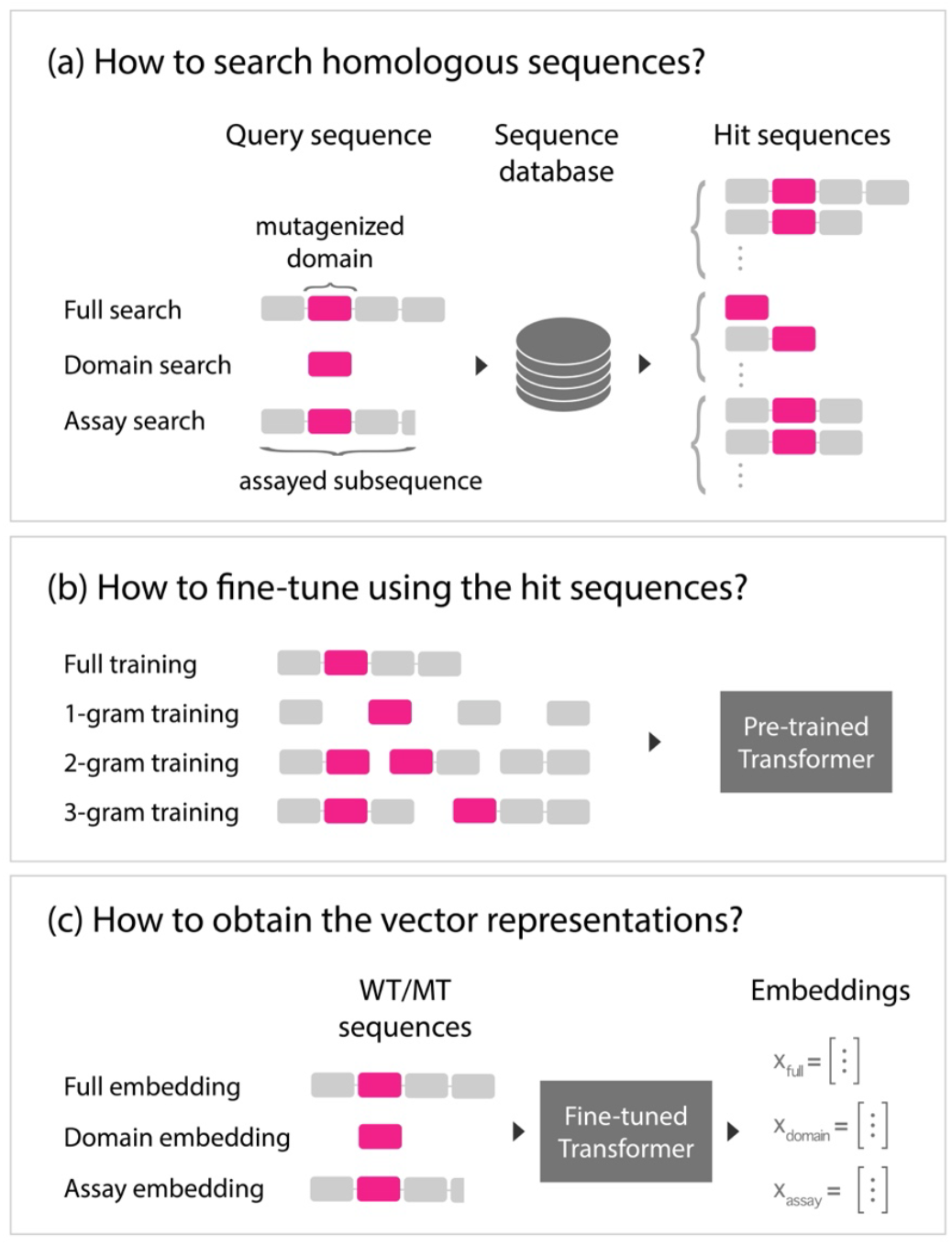
Schematic view of our domain architecture-aware protocols as combinations of homology search, fine-tuning and sequence embedding strategies. See the main text for the detail explanation of each strategy.

a. Homology search. We examined three options on what subsequence to use as a query for homology search. The first option was to use the full-length sequence of a protein (full search); the second one was to extract the mutagenized domain (domain search); the last was to extract the assayed subsequence used in each experiment. As described in Table 1, for some proteins, the assayed subsequences are identical to the full-length sequences or the mutagenized regions; in these cases, the corresponding homology search options become equivalent.
b. Fine-tuning using the hit sequences. We examined whether to use full-length sequences as input to the neural network models (full training). When not using full-length sequences, n-grams (n=1,2,3) were extracted regarding each domain as a unit (n-gram training). For example, in the 2-gram fine-tuning protocol, every two contiguous domain subsequences and the linker region in between was extracted from a full-length sequence. For n-gram generation, we performed domain annotation to each sequence before subsequence extraction. Here the annotation data were only used for n-gram generation, not for any supervision in the experimental pipeline. We provide a schematic explanation on how n-gram training works in Supplementary Figure S1.
c. Vector representations (embeddings) of sequences. Similar to the homology search strategies (a), we tried to use the full length, mutagenized domain, and assayed subsequence of a protein for embedding. Same as explained in (a), the assay embedding coincided with the other embedding options in some cases.

In addition to these major three parts, we also considered several hit sequence filtering methods as post-processing of the sequence generation at the step (b) in our pipeline before fine-tuning: (1) removing sequences not including the mutagenized domain; (2) removing sequences including domains not in a query sequence; (3) the combination of (1) and (2). In Results section, we report the scores with the best performing filtering strategies, while the full results are reported in Supplementary Data S1.

We hypothesized the suitable domain architecture-aware protocol for each protein category from A to C (Table 1) to be as follows (summarized in Table 2, Supplementary Figure S2):

**Table 2.** Summary of the proposed protocol for each protein category. The protocol categories B and C are the ones considering the domain architecture in multi-domain proteins. In the “Embedding” column, assay embedding in parentheses means the described option is equivalent to assay embedding.

| Protocol category | Protein category | Proteins | Homology search | Fine-tuning | Embedding |
| --- | --- | --- | --- | --- | --- |
| A | A | TEM-1, APH(3')-II | Full search | Full length | Full embedding (assay embedding) |
| B | B | Ubiquitin, YAP65, E3 ligase | Domain search | Full length | Domain embedding (assay embedding) |
| C | C | PSD95, Pab1 | Full search | 3-gram | Assay embedding |

A. For the protein category A, in which each protein has a single domain, we propose to combine full search, full training, and full embedding, which were the obvious choices.
B. For the protein category B, which was for the multi-domain proteins without intramolecular domain interactions, we hypothesized that to better incorporate the evolutionary information in the mutagenized domain the appropriate protocol would be the combination of domain search, full training, and domain embedding.
C. For the protein category C, because it was considered crucial to capture the interplay between domains, we selected to perform full search and 3-gram training to cover the interacting sequence parts fully.

On constructing these protocols, we assumed that the best embedding strategy would be the assay embedding because the evolutionary information in the assayed subsequences could directly influence the measured variant effects. Since the assay embedding is equivalent to the full embedding in the category A and domain embedding in the category B, respectively, we employed the corresponding embedding strategies for these categories. We assumed no specific protocols for the protein category D, which was prepared for the proteins not falling into other categories.

We tested these protocol hypotheses by evaluating the variant effect prediction performances on every possible combination of the homology search, fine-tuning, and embedding strategies including the proposed ones above. For comparison, we also set two other protocols: (1) “pre-training” protocol in which the pair of the pre-trained model and full embedding was used without homology search and fine-tuning; (2) “Full” protocol which is the combination of full search only with the length-based filtering (i.e., without filtering based on domain annotation), full training, and full embedding. Note that this protocol was the one evaluated in the previous study [28].

## Results and discussion

### Evotuning protocols substantially improve Transformer-based variant effect prediction accuracy

The evaluation results for the proposed protocols are summarized in Table 3. In almost all cases, the proposed protocols substantially outperformed the respective pre-trained counterparts. This is the first systematically demonstrated results that evotuning can improve Transformer-based variant effect prediction on multi-domain as well as single-domain proteins. We also confirmed that most of the other protocols not described in Table 2 achieved significantly better accuracy than pre-training (Supplementary Data S1). Noticeably, the performance gains were more considerable when the training size was 0.1 compared to 0.8 for most examined proteins, with the maximum improvement of +0.40 in Spearman’s correlation coefficient for YAP65. These results indicate that the protocols effectively incorporate evolutionary information required for accurate variant effect prediction into the learned representations, especially in “low-N” scenarios where the measured values are too scarce to learn evolutionary properties only by supervised learning.

**Table 3.**
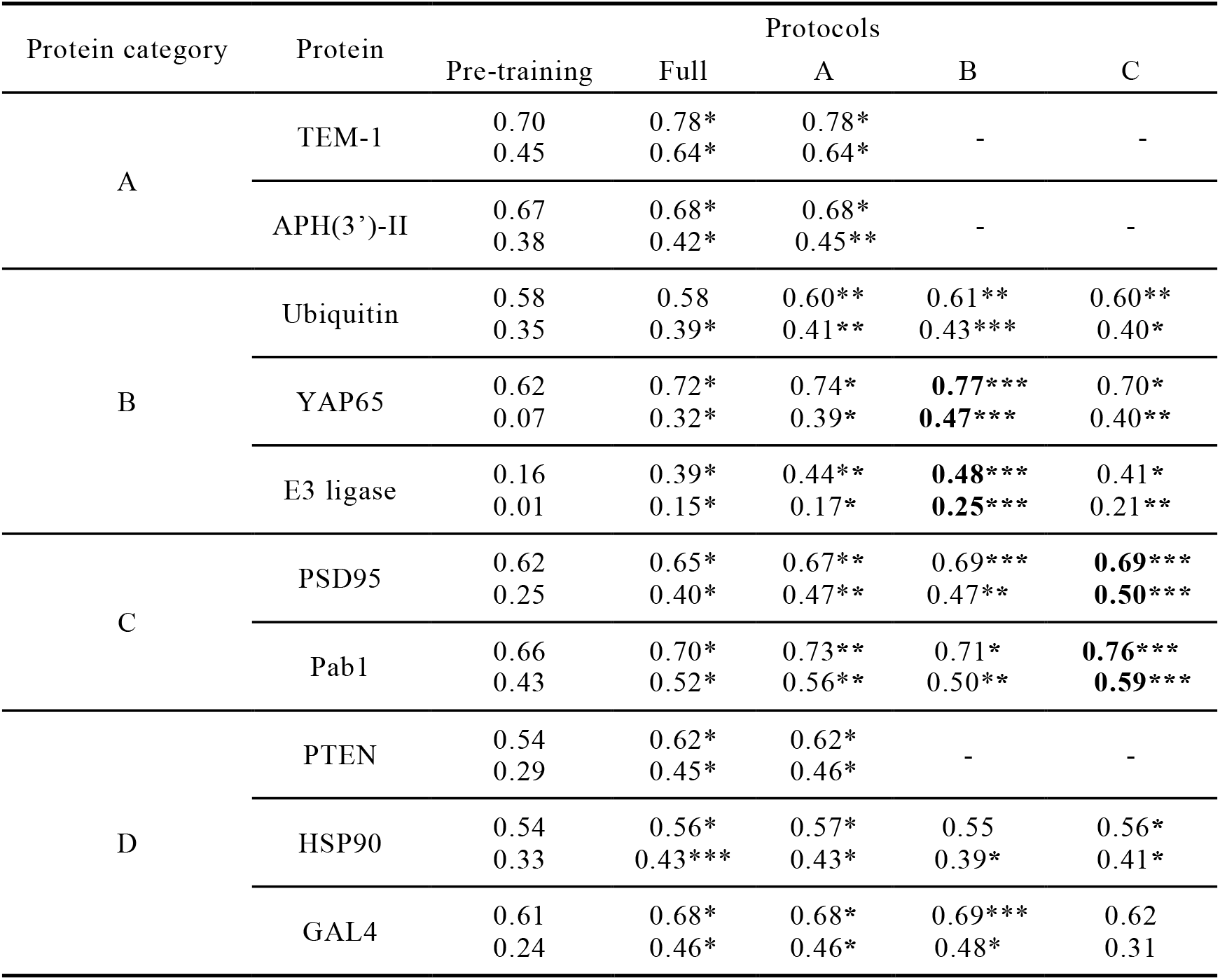
Summary of variant effect prediction accuracy (Spearman’s correlation coefficient). The upper and lower values in each cell are for training size 0.8 and 0.1, respectively. The mean values of Spearman’s correlation coefficients for 32 random seeds are shown. *: significantly (p<0.05; one-sided Wilcoxon test) better than pre-training. **: significantly better than pre-training and protocol “Full”. ***: significantly better than B, C, and either “Full” or A. For the protocols B and C, the best performing protocols are shown in bold (i.e., the scores for both training sizes have *** marks).

We also examined how pre-training affects the variant effect performances by training Transformers from scratch with the same homologous sequences used in the proposed protocols. The resultant prediction accuracy was significantly worse for most cases compared with the pre-trained counterparts (Supplementary Note, Supplementary Table S1).

### Feature analysis: evotuning can incorporate structural information without supervision

An advantage of Transformer is its interpretability provided by attention maps. An attention map quantitatively describes how a residue in a protein affects another residue’s representation, which is internally calculated on embedding. To explore how the substantial performance improvement was realized by evotuning, we visualized the attention maps of YAP65 models as an example. Figure 3 shows the attention maps for the wild-type sequence obtained by the pre-training protocol and the protocol B. We compared these attention maps with the contact map separately computed from the tertiary structure of YAP65. The evotuned model’s attention map displays a contact map-like, roughly symmetric pattern with respect to the diagonal, which is in sharp contrast to the absence of such a pattern for the pre-trained model. This is quite surprising because an attention map is not symmetrized by construction and no explicit knowledge on the tertiary structure was input into the model on fine-tuning. Thus, the visualization suggests that the structural information was implicitly incorporated into the model from the evolutionarily related sequences in an unsupervised manner, possibly leading to the improved variant effect prediction accuracy. A recent study [50] pointed out that a pre-trained Transformer’s attention map can be used for contact map prediction, although the authors explicitly symmetrized the maps before prediction and its implication to variant effect prediction was not discussed.

**Figure 3.**
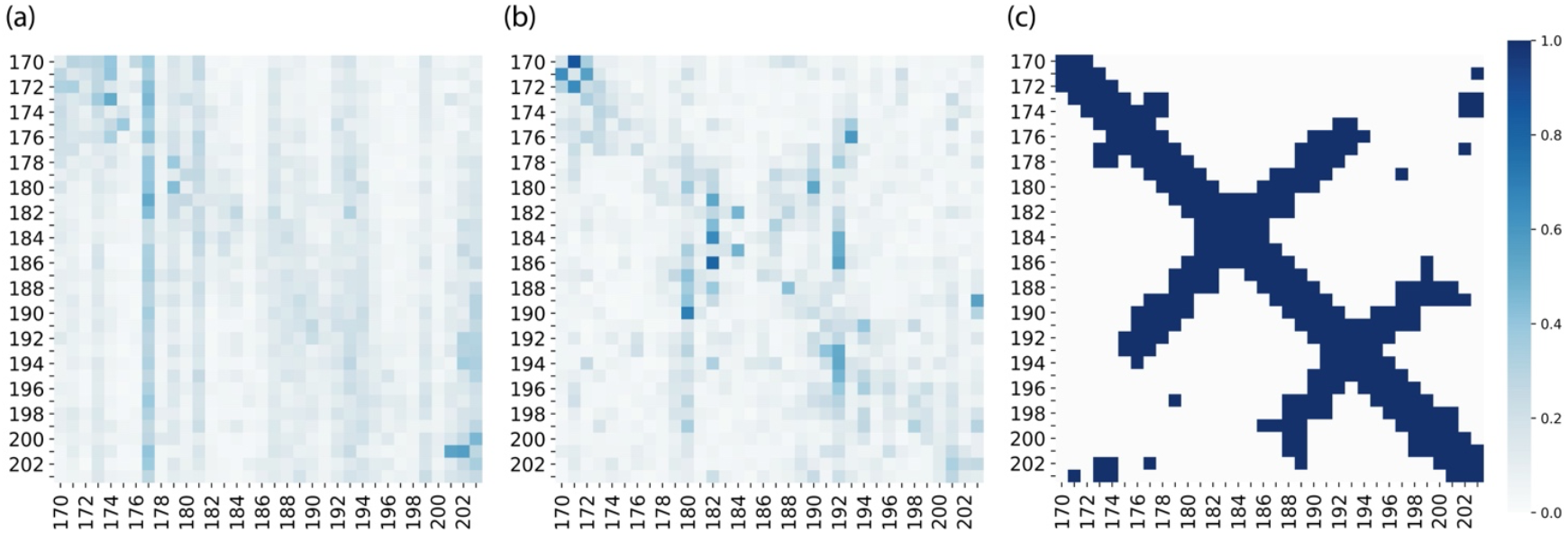
Comparison of YAP65 attention maps obtained by Transformer and the contact map computed from the tertiary structure. (a, b) attention maps obtained by the pre-training protocol (a) and the evotuning protocol B (b). (c) contact map computed from the tertiary structure (PDB: 1JMQ). We calculated the distances between residues (alpha-carbons) with SciPy [49] and regarded a pair of residues within 8 angstroms to be in contact. The mutagenized region (positions from 170 to 203) is visualized. To obtain the attention maps, we computed the max-pooled values over all heads in the last layer. In (a), for comparison with (b), the attention map corresponding to the mutagenized region was extracted from the attention map of the full-length sequence obtained by full embedding, and normalized so that the sum of each row is one.

### Domain architecture-aware protocols achieve the best performances

For single-domain proteins in the category A, the protocols Full and its modified counterpart A improved the prediction accuracies compared to the pre-training (Table 3), which was consistent with the previous study on LSTM [28]. Our results demonstrate that these protocols are also useful for representation learning based on Transformer.

Notably, for the multi-domain proteins in the categories B and C (Table 1), the respective protocols B and C achieved significantly better performances than the protocols Full and A in which evolutionary information was aggregated from the entire sequences across domains. For the proteins in the category B, where one of the multiple independently working domains was mutagenized, the protocol exploiting the mutagenized domain sequence only (i.e., protocol B) achieved the best scores: they were either the largest among all the protocols or at least larger than the protocol Full. By contrast, when the protocol C was applied to this protein category, several cases were observed to be merely comparable to the protocol Full (Ubiquitin with training size 0.1, YAP65 with 0.1, and E3 ligase with 0.8). These results indicate that the careful treatment of the domain architecture and the intramolecular domain interactions indeed lead to improved protein representations.

Next, for the category C proteins with cooperative domains, the protocol aiming to gather evolutionary information from sequences covering inter-domain interactions (i.e., protocol C) realized the best results. In a rare case (PSD95 with training size 0.8), the performance of the protocol B was comparable to the protocol C. A possible reason is that the mutagenesis assay was conducted for the isolated PDZ domain, not for the full length [16]. Nonetheless, in the remaining cases, the protocol C was better than the protocol B. The result again demonstrates the validity of our hypothesis that the representation learning of multi-domain proteins can be improved when explicitly considering different domain architectures of proteins. We also tried other training sizes (0.9, 0.5, 0.3, and 0.01) and found consistent results (Supplementary Data S2).

Lastly, no common superior protocol was confirmed for the protein category D in contrast to A, B, and C. Because the mutations were introduced to the whole sequence of PTEN [9], we only conducted the protocols Full and A experiments not considering domain architectures, which resulted in the improved performances compared with the pre-training. This is consistent with the results for the protocols Full and A applied to the other proteins, indicating the partial effectiveness of evotuning. For HSP90 with training size 0.1, the protocols Full and A performed the best while it was comparable to the protocol C with training size 0.8. This performance mismatch between training sizes was also seen for GAL4 with the protocol B but not observed in the other proteins. We speculate that the difference stems from the distinct working mechanisms for these HSP90 and GAL4 (necessities for homodimer formation) from the other proteins.

To assess our approach more thoroughly, we also examined other algorithms for variant effect prediction (Supplementary Note). Compared with the previous LSTM-based approach [28], our Transformer-based protocols were shown to be significantly more accurate for most cases (Supplementary Table S2). Also, when compared with a weakly-supervised algorithm from DeepSequence [51], our LASSO-LARS predictors on top of the evotuned models supervised by the measured variant effects were demonstrated to outperform when training size was 0.8 (Supplementary Tables S3-S4). On the other hand, with training size 0.1, DeepSequence was better than our models for several proteins. As another assessment, we examined how end-to-end training, instead of first freezing the evotuned weights and then training LASSO-LARS as described, can affect the prediction performances. The evaluation results in Supplementary Table S5 indicates that the accuracies were greatly improved for most proteins with training size 0.8; however, LASSO-LARS performed consistently better with training size 0.1. Overall, these results indicate the effectiveness of our domain architecture-aware protocols; at the same time, it is also suggested that we still have rooms for algorithmic improvement in “low-N” regime to exploit measured variant effects further.

### Effectiveness of domain architecture-aware sequence filtering for evotuning

The exhaustive evaluation verified that our domain architecture-aware sequence filtering strategies effectively improved prediction accuracy: by filtering, comparable or superior performances were achieved for all proteins examined (Supplementary Data S1). Our filtering method has the novelty in that it is designed for multi-domain proteins considering their domain architectures and mutagenized domains (Materials and methods). Recently, in a study [52], sequence filtering based on Levenshtein distance was performed to evotune LSTM-based representation models for variant effect prediction. However, it is unclear whether it can be successfully applied to multi-domain proteins because the study’s evaluations were only performed for single-domain proteins (GFP and TEM-1).

## Conclusion

In this work, we proposed the domain architecture-aware protocols for Transformer representation learning of multi-domain proteins, combining homology search, fine-tuning, and sequence vectorization strategies. By systematically evaluating the variant effect prediction performances on ten different protein engineering tasks, we confirmed the effectiveness of our protocols: 1) they gave substantial performance gains compared with the pre-training, especially in “low-N” scenarios where the measured values are scarce; 2) the best-performing protocols were constructed by adequately considering the domain architecture and intramolecular domain interactions of the engineering target protein.

While the evotuning protocols were demonstrated to be crucial for variant effect prediction, some protein engineering scenarios were not covered in this study. One example is protein engineering focusing on linker regions between domains [53], for which the protocols B and C cannot be directly applied. Another example is intramolecular interactions between distal domains (e.g., the interaction between N-terminus and C-terminus far apart by multiple domains), which can be regarded as an extension of the protein category C. In addition, potentially large regions of linker or unannotated sequence may pose another issue. The optimal evotuning protocols for these cases should be addressed in a future study. We also note that although the proposed protocols (Table 2) achieved better accuracy than the “Full” protocol, there were other protocols that showed comparable accuracy to the proposed protocols (Supplementary Data S1). These included, for example, a variation of the protocol B using 1-gram training instead of full training, which corresponds to extracting the mutagenized domain from homologous sequences. The reason for performance differences observed between these protocols could not be clarified in this study. Lastly, because the base model in this study was pre-trained with Pfam containing single domain sequences, this could negatively impact variant effect prediction, especially for multi-domain proteins. Starting with other large-scale databases composed of whole protein sequences such as UniRef for evotuning would be another point for further improvement.

## Key points

- In machine learning-guided protein engineering, it is crucial to incorporate evolutionary information specific to a target protein into the feature vectors. However, no protocols are established for Transformer-based variant effect prediction, in which features are learned and generated from unlabeled homologous sequences. To this end, this study proposes effective protocols combining homology search, fine-tuning, and sequence vectorization strategies for multi-domain proteins as well as single-domain ones.
- By systematic performance evaluation on various tasks, we demonstrated the protein representations obtained by the proposed protocols significantly outperformed the ones by pre-training as well as a previous LSTM-based approach for most cases. In particular, in “low-N” scenarios where the number of labeled training data available is small, the proposed protocols were shown to give substantial performance gains; at the same time, we found that we still have rooms for developing more data-efficient algorithms in this regime.
- We also confirmed that the best-performing protocols were constructed by adequately considering the engineered protein’s domain architecture and intramolecular domain interactions. The visualization of the attention maps in Transformer given by the best protocol suggested that structural information was implicitly captured by the model, which was a possible reason for the performance improvement. These results indicate that the proposed protocols are effective and indispensable for protein engineering.

## Data availability

The data underlying this article are available in the article and in its online supplementary material. The source codes are available at https://github.com/dlnp2/evotuning_protocols_for_transformers.

## Author contributions statement

H.Y. conceived of the study, developed the method, wrote the code, analyzed the data, and wrote the paper. Y.S. coordinated the project, participated in the data interpretation, and wrote the paper.

## Acknowledgements

For neural network training, computational resource of AI Bridging Cloud Infrastructure (ABCI) provided by National Institute of Advanced Industrial Science and Technology (AIST) was used. Computations were partially performed on the NIG supercomputer at ROIS National Institute of Genetics.

---

**Supplementary Figure S1.**
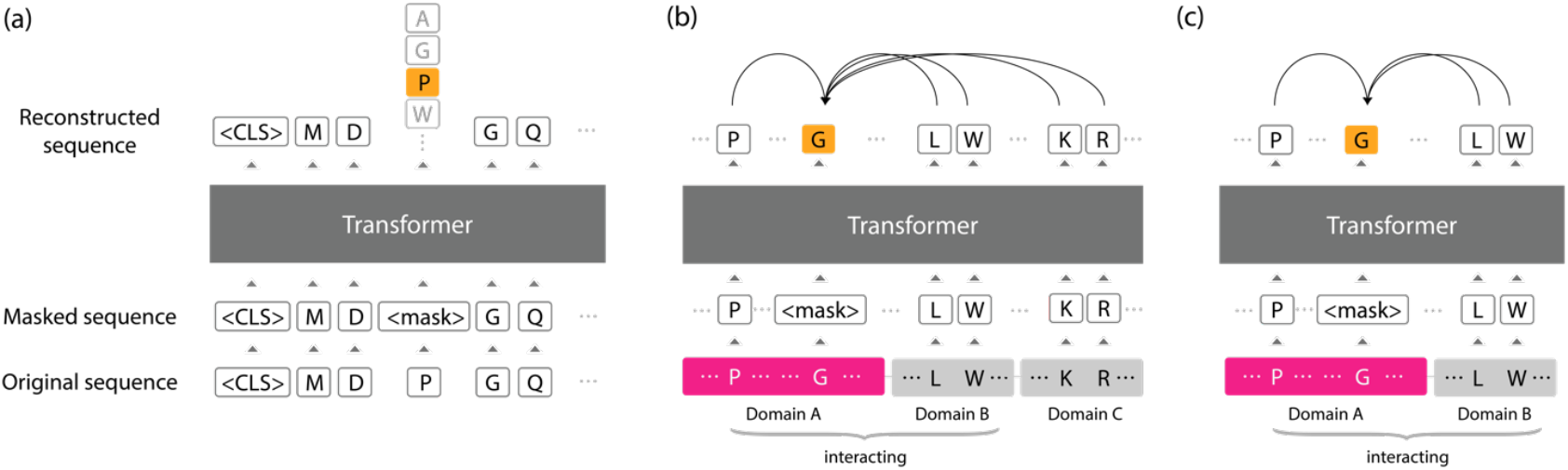
(a) Masked language modeling task in this study. An original protein sequence is first corrupted to include some “<mask>” tokens instead of the original amino acids according to a given probability (“P” is replaced in this figure). Then, the corrupted sequence is fed into a Transformer model for training to predict the original amino acids at the “<mask>” positions, using information from other amino acids. (b, c) Schematic explanation of the proposed protocols. We consider a wild-type protein with three domains named A, B, and C in these panels. Here we assume that (1) the domain A is the mutagenesis target (highlighted in pink) and (2) the domain A and B have some physically interacting residues between them, which would affect how the former works. Our goal in masked language modeling is to obtain a high-quality representation of the domains of interest. (b) Schematic explanation of “Full” evotuning protocol. In this approach, we try to restore the corrupted token (“G”) based on every other residue in the original full-length sequence. However, information from the domain C probably serves as noise because it is not relevant to the interaction between A and B. (c) Schematic explanation of “n-gram” evotuning protocol (in this case “2-gram”). In short, n-gram training means fine-tuning a model on all n-grams over domains containing the mutagenized domain in masked language modeling. In contrast to “Full” protocol, we focus on the interacting, contiguous domains (A and B) by removing the domain C on training. Since in this study we regard each domain as a “word”, or unigram, as in NLP, this approach is named “2-gram”. Here, we also include the linker region between the domains A and B, which is usually considered short enough compared to the domains themselves.

**Supplementary Figure S2.**
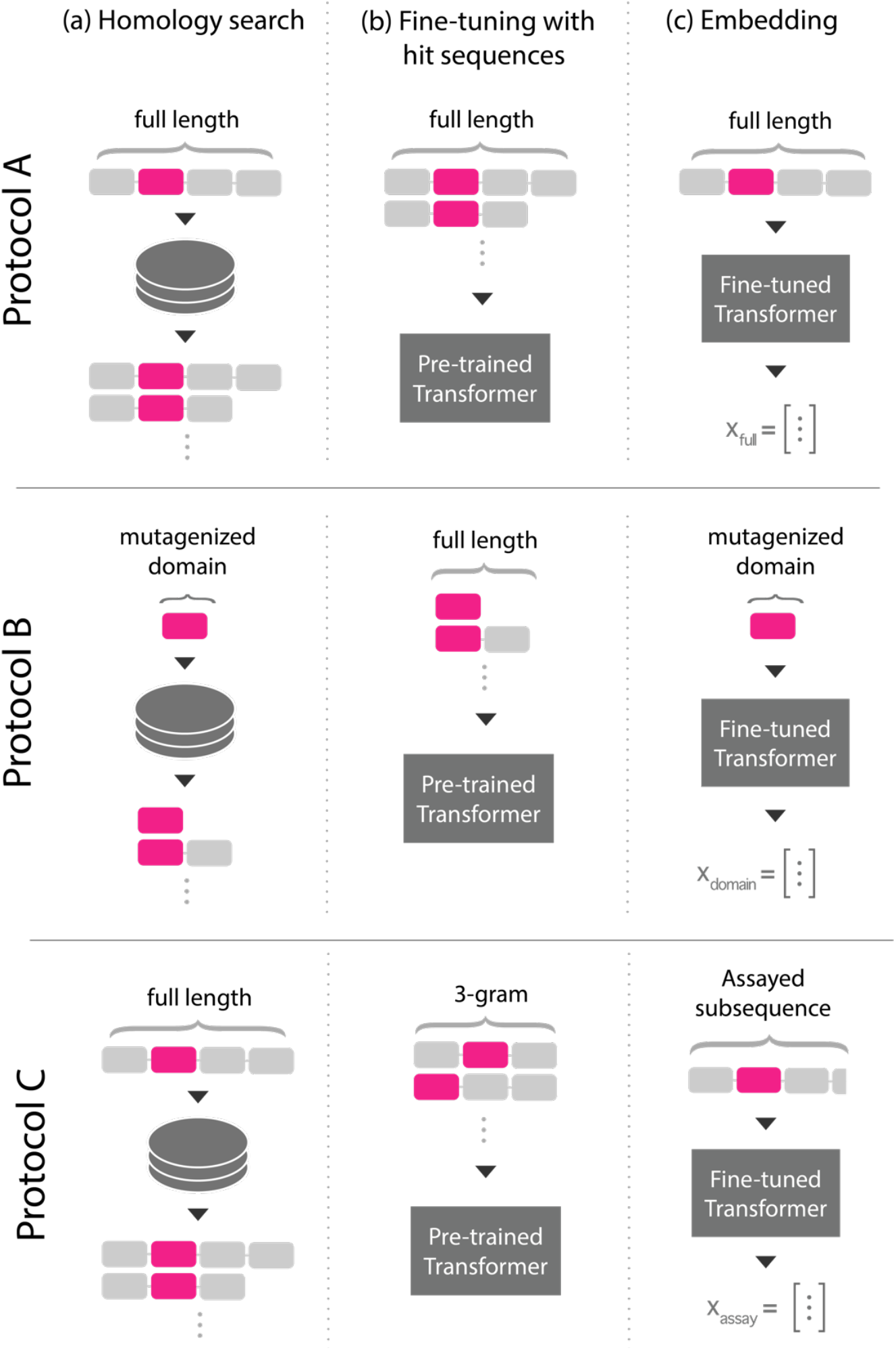
Schematic summary of the proposed protocols (described in Table 2). The protocols A, B, and C differ from each other in the domain architectures of sequences used for (a) homology search, (b) fine-tuning the pre-trained Transformer, and (c) embedding through the fine-tuned Transformer. Each rectangle in a sequence indicates a domain. The mutagenized domains are highlighted in pink.

## Supplementary Note

### Comparison of variant effect prediction accuracy between models with and without pre-training

To evaluate how pre-training Transformer-based protein representation models affects variant effect prediction accuracy, we compared the performances with and without pre-training for each protein. The only difference between these two approaches is whether we include the step (a) in our experimental pipeline described in Figure 1: when we skip pre-training, the whole neural network was initialized with random parameters. As Supplementary Table S1 demonstrates, the pre-trained models consistently outperformed the models without pre-training, with large margins especially for training size = 0.1. This result indicates that it is crucial to incorporate the general knowledge on proteins into the Transformer-based models before fine-tuning for them to specialize in a target protein toward accurate variant effect prediction.

### Comparison of variant effect prediction accuracy between our Transformer-based and LSTM-based evotuned models

To assess the effectiveness of our protocols, we compared the variant effect prediction accuracy against the LSTM-based approach from UniRep [28]. Because the original TensorFlow implementation of UniRep is highly computational resource demanding, we instead used a more efficient JAX [54] implementation [55]. For evaluation for each protein, we initialized the LSTM model with the same pre-trained weight as the original one and trained according to our “Full” protocol to emulate the original experiments in [28] (refer for the definition of “Full” protocol to Table 2), with learning rate 0.00001, 65 epochs, and batch size selected from 32, 64, 128 to fit into a single NVIDIA V100 GPU memory. As summarized in Supplementary Table S2, our Transformer-based models significantly outperformed LSTM-based models for most cases, which demonstrates the effectiveness of our protocols.

### Comparison of variant effect prediction accuracy between our supervised models and weakly-supervised models

To further assess the effectiveness of our protocols, we compared the variant effect prediction accuracy against a weakly-supervised approach from DeepSequence [51], which is based on variational autoencoders (VAEs) [53]. In short, for training, a DeepSequence model requires a multiple sequence alignment (MSA) composed of homologous sequence fragments obtained by jackhmmer using the target protein’s mutagenized domain sequence as the query. The MSA is post-processed not to include positions with more than 30% gaps nor sequence fragments with more than 50% gaps. For prediction, the difference between the log likelihoods of a mutagenized sequence and a wild-type sequence is used as a surrogate indicator of variant effect. As such, DeepSequence does not need a downstream supervised predictor, which is distinguished from our supervised approach that uses LASSO-LARS. It is worth mentioning that another requirement for DeepSequence stemming from its network architecture is the length of the query sequence for homology search and that of sequences used for prediction (i.e., embedding) is the same.

For maximally fair performance comparison, we evaluated DeepSequence with the proposed protocols described in Table 2: we used the MSAs with our own homologs obtained by the protocols and the same embedding strategies. However, since DeepSequence has the requirement of the same sequence length as described above, it is impossible to evaluate DeepSequence with the protocol C, which uses the different lengths of sequences in homology search and embedding (the full-length sequence in homology search and the assay sequence in embedding; Table 2). Thus, for the proteins in category C, PSD95 and Pab1, we applied the protocol B as considered to be done in the original work [51]. For training, we set the default hyperparameters.

The evaluation results are summarized in Supplementary Table S3. Our models achieved much better accuracy than DeepSequence with training size = 0.8. However, for training size = 0.1, DeepSequence was generally better than our models. To ensure that our evaluation procedure was correct, we compared the results with the reported results from [51] summarized in Supplementary Table S4 and confirmed these two were relatively similar. The discrepancies observed are considered to be derived from the differences between homology search databases: the original DeepSequence used UniRef100, whereas ours was UniRef50, which is far smaller. These evaluation results suggest that we still have rooms for algorithmic improvement in “low-N” regime to exploit measured variant effects for supervision further.

### Evaluation of end-to-end variant effect predictors

As another evaluation of our protocols, we performed variant effect prediction by end-to-end training rather than LASSO-LARS. Specifically, for each protein, an evotuned Transformer attached with a single fully connected layer on top of it was trained to regress the measured variant effects. The embeddings for each amino acid were averaged to produce a fixed-length vector fed into the top layer. The learning rate was 0.00001 and the batch size was 16.

Supplementary Table S5 is the evaluation summary. For training size = 0.8, the performances were greatly improved by end-to-end training compared with LASSO-LARS based approach, with the noticeable exceptions of YAP65 and E3 ligase. The possible reason is the relatively small numbers of variants for these two proteins that have only an order of magnitude fewer variants than other proteins (Table 1). In contrast, for training size = 0.1, the performances were generally better for LASSO-LARS based models. These results suggest that the evotuned models had successfully exploited the rich measurement data in “high-N” regime but may have benefitted from the LASSO’s ability to select sparse features specifically in the “low-N” regime to avoid over-fitting.

**Supplementary Table S1.**
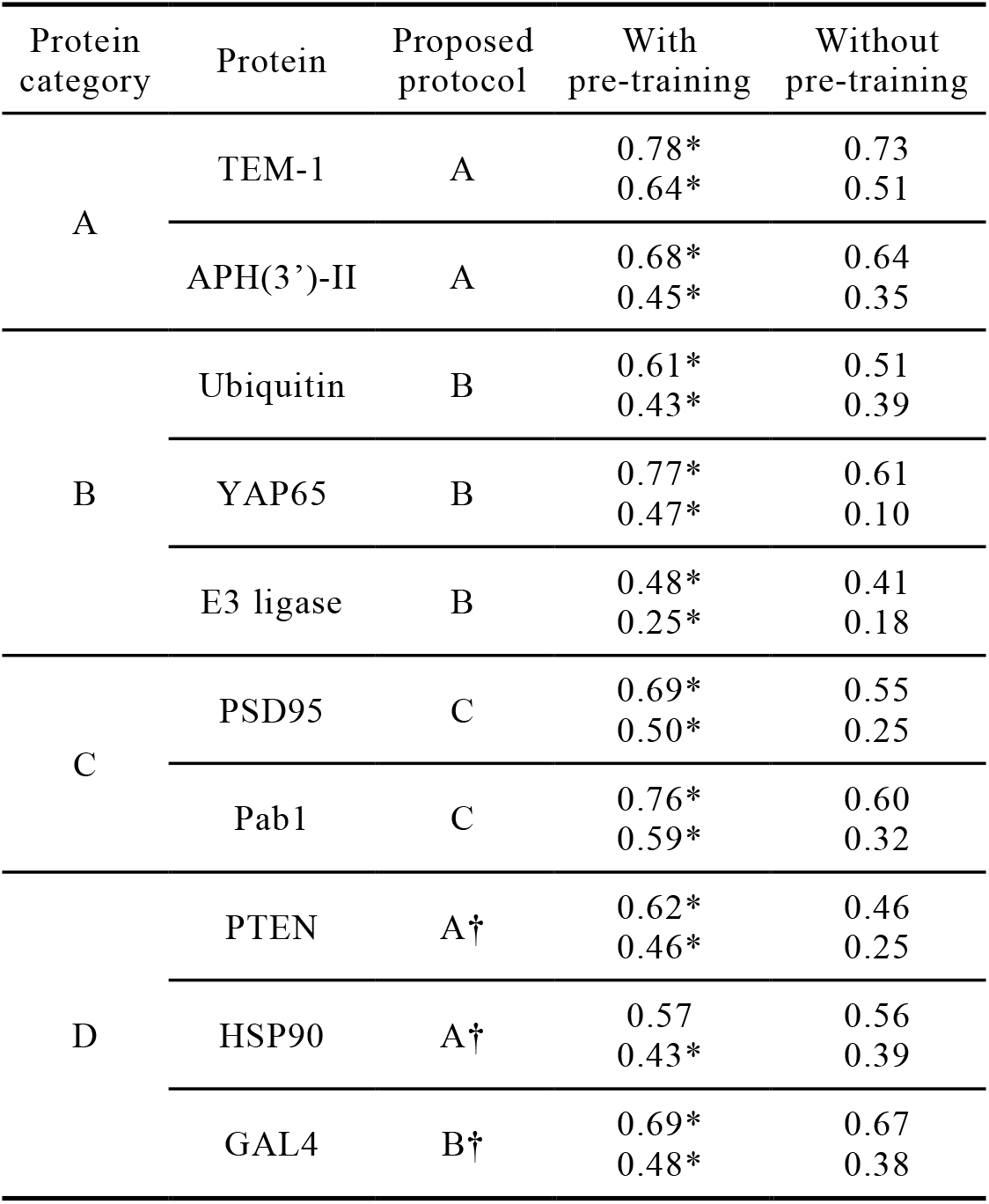
Comparison of variant effect prediction accuracy (Spearman’s correlation coefficient) between models with and without pre-training, which were subsequently fine-tuned with respective homologous sequences obtained by the proposed protocols. The upper and lower values in each cell are for training size 0.8 and 0.1, respectively. Refer to Table 2 for the definitions of the proposed protocols. *: significantly (p<0.05; one-sided Wilcoxon test) better than “without pre-training” setting. †: best-performing protocols for each protein in the protein category D.

**Supplementary Table S2.**
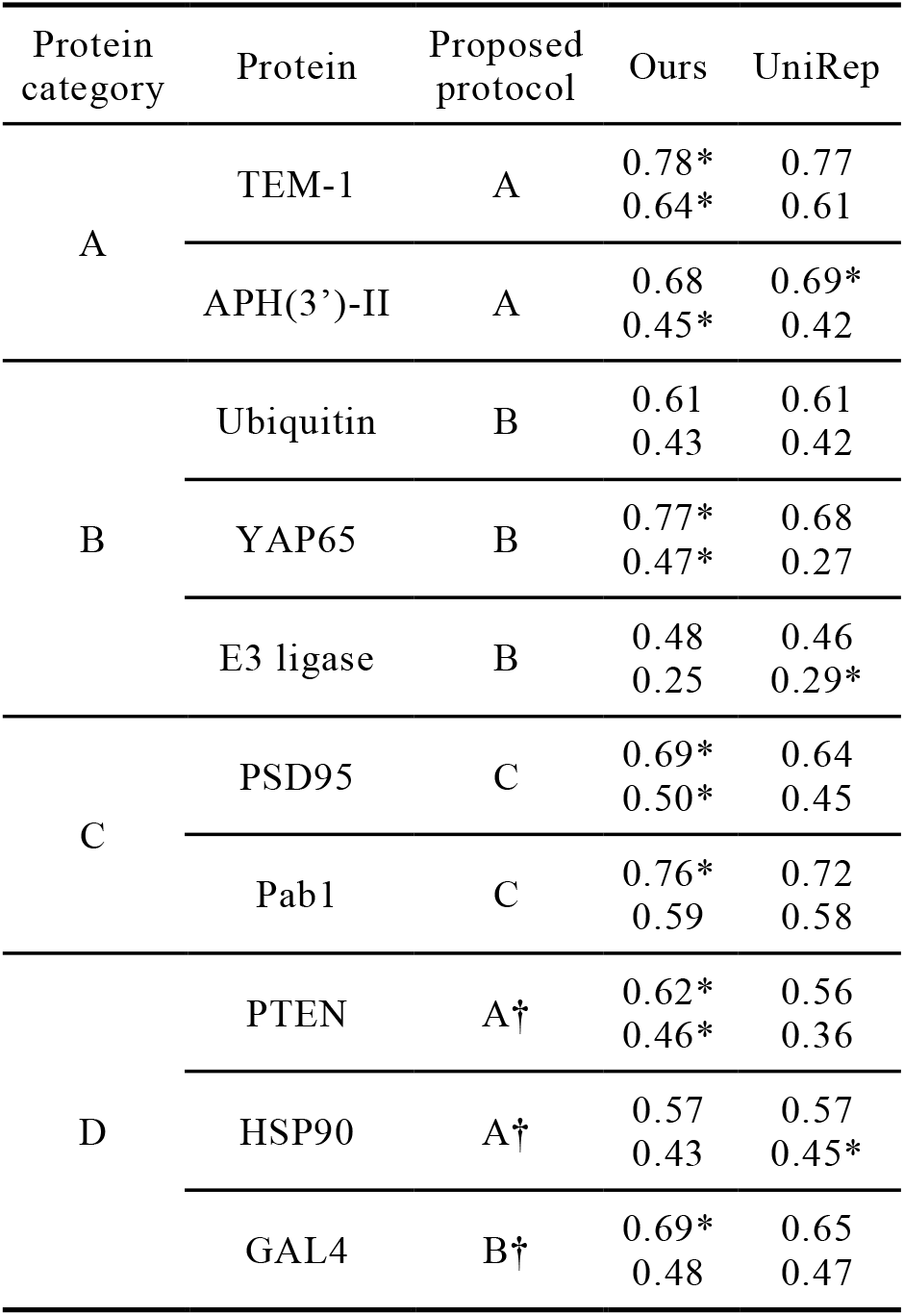
Comparison of variant effect prediction accuracy (Spearman’s correlation coefficient) between Transformer-based (“Ours”) and LSTM-based (“UniRep”) evotuned models. The upper and lower values in each cell are for training size 0.8 and 0.1, respectively. Refer to Table 2 for the definitions of the proposed protocols. *: significantly (p<0.05; one-sided Wilcoxon test) better than the other model. †: best-performing protocols for each protein in the protein category D.

**Supplementary Table S3.** Comparison of variant effect prediction accuracy (Spearman’s correlation coefficient) between our supervised models and weakly-supervised models from DeepSequence trained with our own homologs. The upper and lower values in each cell in “Ours” column are for training size 0.8 and 0.1, respectively. For APH(3’)-II, PTEN, and HSP90, we did not perform evaluation on DeepSequence because of the lack of training homolog sequences after alignment filtering as performed in [51] (denoted as “-”). Refer to Table 2 for the definitions of the proposed protocols. †: best-performing protocols for each protein in the protein category D.

| Protein category | Protein | Proposed protocol | Ours | DeepSequence (ours) |
| --- | --- | --- | --- | --- |
| A | TEM-1 | A | 0.78<br>0.64 | 0.71 |
|  | APH(3')-II | A | 0.68<br>0.45 | - |
| B | Ubiquitin | B | 0.61<br>0.43 | 0.35 |
|  | YAP65 | B | 0.77<br>0.47 | 0.62 |
|  | E3 ligase | B | 0.48<br>0.25 | 0.48 |
| C | PSD95 | C | 0.69<br>0.50 | 0.57 |
|  | Pab1 | C | 0.76<br>0.59 | 0.65 |
| D | PTEN | A† | 0.62<br>0.46 | - |
|  | HSP90 | A† | 0.57<br>0.43 | - |
|  | GAL4 | B† | 0.69<br>0.48 | 0.43 |

**Supplementary Table S4.** Comparison of variant effect prediction accuracy (Spearman’s correlation coefficient) between our supervised models and weakly-supervised models from DeepSequence. The upper and lower values in each cell in “Ours” column are for training size 0.8 and 0.1, respectively. The values in “DeepSequence” are from [51]. The value for YAP65 was not reported in [51] (thus, denoted as “-”). Refer to Table 2 for the definitions of the proposed protocols. †: best-performing protocols for each protein in the protein category D.

| Protein category | Protein | Proposed protocol | Ours | DeepSequence [51] |
| --- | --- | --- | --- | --- |
| A | TEM-1 | A | 0.78<br>0.64 | 0.78 |
|  | APH(3')-II | A | 0.68<br>0.45 | 0.66 |
| B | Ubiquitin | B | 0.61<br>0.43 | 0.42 |
|  | YAP65 | B | 0.77<br>0.47 | - |
|  | E3 ligase | B | 0.48<br>0.25 | 0.54 |
| C | PSD95 | C | 0.69<br>0.50 | 0.60 |
|  | Pab1 | C | 0.76<br>0.59 | 0.69 |
| D | PTEN | A† | 0.62<br>0.46 | 0.42 |
|  | HSP90 | A† | 0.57<br>0.43 | 0.53 |
|  | GAL4 | B† | 0.69<br>0.48 | 0.49 |

**Supplementary Table S5.**
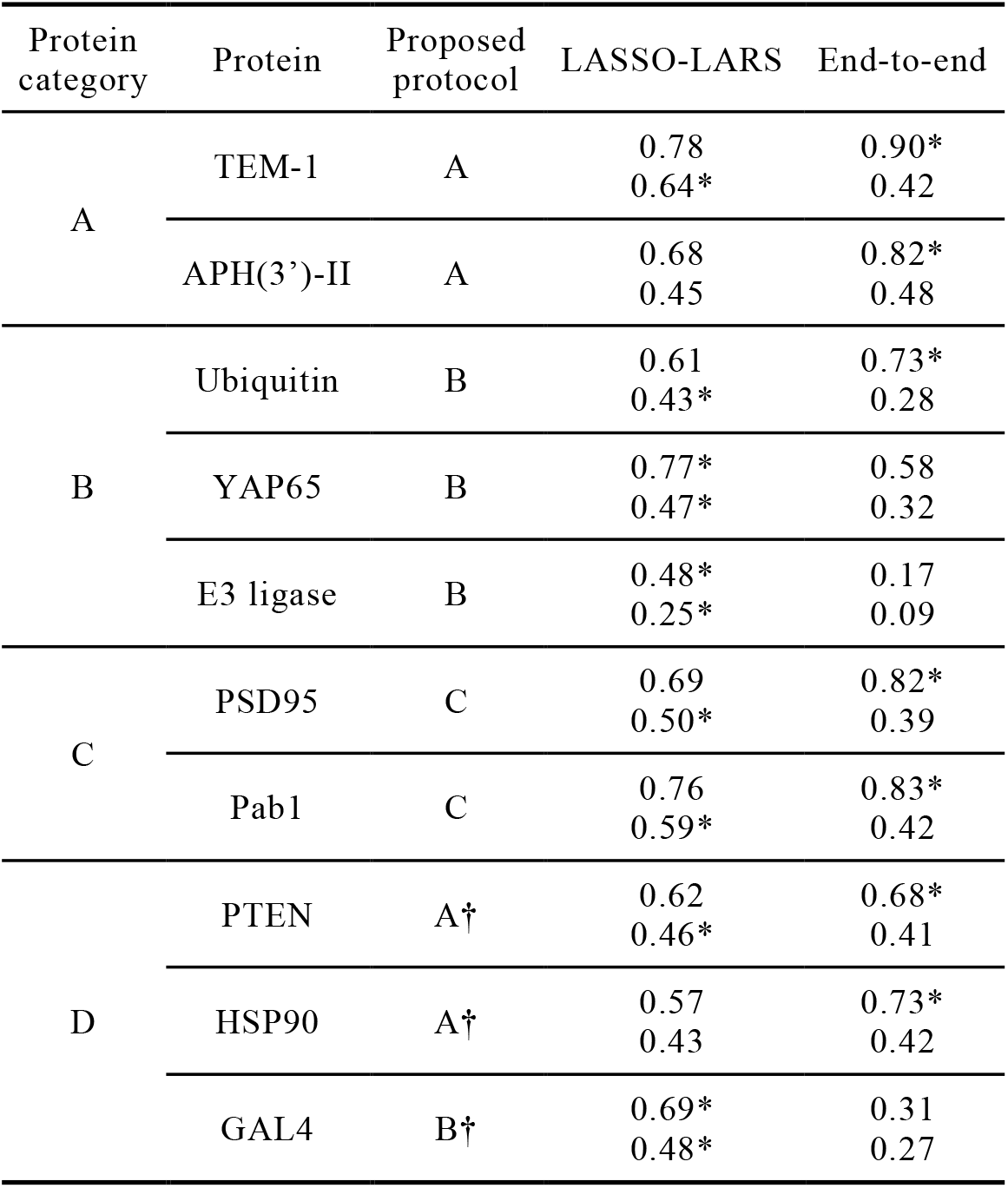
Comparison of variant effect prediction accuracy (Spearman’s correlation coefficient) between two types of predictors: in “LASSO-LARS” column, we froze an evotuned model weight and trained a LASSO-LARS predictor on top of it; in “End-to-end” column, we attached a fully connected layer on an evotuned Transformer and trained the whole network for variant effect prediction. The upper and lower values in each cell are for training size 0.8 and 0.1, respectively. Refer to Table 2 for the definitions of the proposed protocols. †: best-performing protocols for each protein in the protein category D.

## References

[1] Chen K, Arnold FH. Tuning the activity of an enzyme for unusual environments: sequential random mutagenesis of subtilisin E for catalysis in dimethylformamide. Proc Natl Acad Sci U S A. 1993;90:5618–22.

[2] Pédelacq JD, Cabantous S, Tran T et al. Engineering and characterization of a superfolder green fluorescent protein. Nat Biotechnol. 2006;24:79–88.

[3] Levin AM, Bates DL, Ring AM et al. Exploiting a natural conformational switch to engineer an interleukin-2 ‘superkine’. Nature. 2012;484:529–33.

[4] Gaudelli NM, Komor AC, Rees HA et al. Programmable base editing of A•T to G•C in genomic DNA without DNA cleavage. Nature. 2017;551:464–471.

[5] Ekman D, Björklund AK, Frey-Skött J et al. Multi-domain proteins in the three kingdoms of life: orphan domains and other unassigned regions. J Mol Biol. 2005;348:231–43.

[6] Ahmad ZA, Yeap SK, Ali AM et al. scFv antibody: principles and clinical application. Clin Dev Immunol. 2012;2012:980250.

[7] Makarova KS, Wolf YI, Iranzo et al. Evolutionary classification of CRISPR-Cas systems: a burst of class 2 and derived variants. Nat Rev Microbiol. 2020;18:67–83.

[8] Gray VE, Hause RJ, Luebeck J et al. Quantitative Missense Variant Effect Prediction Using Large-Scale Mutagenesis Data. Cell Syst. 2018;6:116–124.e3.

[9] Matreyek KA, Starita LM, Stephany JJ et al. Multiplex assessment of protein variant abundance by massively parallel sequencing. Nat Genet. 2018;50:874–882.

[10] Kitzman JO, Starita LM, Lo RS et al. Massively parallel single-amino-acid mutagenesis. Nat Methods. 2015;12:203–6.

[11] Firnberg E, Labonte JW, Gray JJ et al. A comprehensive, high-resolution map of a gene’s fitness landscape. Mol Biol Evol. 2014;31:1581–92.

[12] Melnikov A, Rogov P, Wang L et al. Comprehensive mutational scanning of a kinase in vivo reveals substrate-dependent fitness landscapes. Nucleic Acids Res. 2014;42:e112.

[13] Roscoe BP, Bolon DN. Systematic exploration of ubiquitin sequence, E1 activation efficiency, and experimental fitness in yeast. J Mol Biol. 2014;426:2854–70.

[14] Fowler DM, Araya CL, Fleishman SJ et al. High-resolution mapping of protein sequence-function relationships. Nat Methods. 2010;7:741–6.

[15] Starita LM, Pruneda JN, Lo RS et al. Activity-enhancing mutations in an E3 ubiquitin ligase identified by high-throughput mutagenesis. Proc Natl Acad Sci U S A. 2013;110:E1263–72.

[16] McLaughlin RN Jr, Poelwijk FJ, Raman A et al. The spatial architecture of protein function and adaptation. Nature. 2012;491:138–42.

[17] Melamed D, Young DL, Gamble CE et al. Deep mutational scanning of an RRM domain of the Saccharomyces cerevisiae poly(A)-binding protein. RNA. 2013;19:1537–51.

[18] Saito Y, Oikawa M, Nakazawa H et al. Machine-Learning-Guided Mutagenesis for Directed Evolution of Fluorescent Proteins. ACS Synth Biol. 2018;7:2014–2022.

[19] Wu Z, Kan SBJ, Lewis RD et al. Machine learning-assisted directed protein evolution with combinatorial libraries. Proc Natl Acad Sci U S A. 2019;116:8852–8858.

[20] Bedbrook CN, Yang KK, Robinson JE et al. Machine learning-guided channelrhodopsin engineering enables minimally invasive optogenetics. Nat Methods. 2019;16:1176–1184.

[21] Rao R, Bhattacharya N, Thomas N et al. Evaluating Protein Transfer Learning with TAPE. In: 33rd Conference on Neural Information Processing Systems (NeurIPS 2019), Vancouver, Canada.

[22] Elnaggar A, Heinzinger M, Dallago C et al. ProtTrans: Towards Cracking the Language of Life’s Code Through Self-Supervised Deep Learning and High Performance Computing. bioRxiv doi: 10.1101/2020.07.12.199554.

[23] Rives A, Meier J, Sercu T et al. Biological structure and function emerge from scaling unsupervised learning to 250 million protein sequences. bioRxiv doi: 10.1101/622803.

[24] Kawashima S, Kanehisa M. AAindex: amino acid index database. Nucleic Acids Res. 2000;28:374.

[25] Tian F, Zhou P, Li Z. T-scale as a novel vector of topological descriptors for amino acids and its application in QSARs of peptides. J Mol Struct. 2007;830:106–115.

[26] Yang KK, Wu Z, Bedbrook CN et al. Learned protein embeddings for machine learning. Bioinformatics. 2018;34:2642–2648.

[27] Krause B, Lu L, Murray I et al. Multiplicative LSTM for sequence modelling. arXiv e-prints, page arXiv:1609.07959, September 2016.

[28] Alley EC, Khimulya G, Biswas S et al. Unified rational protein engineering with sequence-based deep representation learning. Nat Methods. 2019;16:1315–1322.

[29] Vaswani A, Shazeer N, Parmar N et al. Attention is all you need. In Advances in Neural Information Processing Systems, 2017, 6000–6010.

[30] Wang A, Singh A, Michael J et al. Glue: A multi-task benchmark and analysis platform for natural language understanding. In Proceedings of the 2018 EMNLP Workshop BlackboxNLP: Analyzing and Interpreting Neural Networks for NLP, pages 353–355.

[31] Wang A, Pruksachatkun Y, Nangia N et al. SuperGLUE: A Stickier Benchmark for General-Purpose Language Understanding Systems. In: 33rd Conference on Neural Information Processing Systems (NeurIPS 2019), Vancouver, Canada.

[32] El-Gebali S, Mistry J, Bateman A, Eddy SR et al. The Pfam protein families database in 2019. Nucleic Acids Res. 2019;47:D427–D432.

[33] Paszke A, Gross S, Massa F et al. PyTorch: An Imperative Style, High-Performance Deep Learning Library. In Advances in Neural Information Processing Systems, 2019, 8024–8035.

[34] Kingma DP and Ba J. Adam: A Method for Stochastic Optimization. arXiv e-prints, page arXiv:1412.6980, December 2014.

[35] Micikevicius P, Narang S, Alben J et al. Mixed Precision Training. arXiv e-prints, page arXiv:1710.03740, October 2017.

[36] Devlin J, Chang M-W, Lee K et al. BERT: Pre-training of Deep Bidirectional Transformers for Language Understanding. In: Proceedings of the 2019 Conference of the North American Chapter of the Association for Computational Linguistics: Human Language Technologies, Volume 1 (Long and Short Papers)

[37] Eddy SR. Accelerated Profile HMM Searches. PLoS Comput Biol. 2011;7:e1002195.

[38] Suzek BE, Wang Y, Huang H et al. UniRef clusters: a comprehensive and scalable alternative for improving sequence similarity searches. Bioinformatics. 2015;31:926–32.

[39] Yu L, Tanwar DK, Penha EDS et al. Grammar of protein domain architectures. Proc Natl Acad Sci U S A. 2019;116:3636–3645.

[40] Laursen L, Kliche J, Gianni S et al. Supertertiary protein structure affects an allosteric network. Proc Natl Acad Sci U S A. 2020;117:24294–24304.

[41] Deo RC, Bonanno JB, Sonenberg N et al. Recognition of polyadenylate RNA by the poly(A)-binding protein. Cell. 1999;98:835–45.

[42] Safaee N, Kozlov G, Noronha AM et al. Interdomain allostery promotes assembly of the poly(A) mRNA complex with PABP and eIF4G. Mol Cell. 2012;48:375–86.

[43] Lee JO, Yang H, Georgescu MM et al. Crystal structure of the PTEN tumor suppressor: implications for its phosphoinositide phosphatase activity and membrane association. Cell. 1999;99:323–34.

[44] Mishra P, Flynn JM, Starr TN et al. Systematic Mutant Analyses Elucidate General and Client-Specific Aspects of Hsp90 Function. Cell Rep. 2016;15:588–598.

[45] Richter K, Muschler P, Hainzl O, et al. Coordinated ATP hydrolysis by the Hsp90 dimer. J Biol Chem. 2001 Sep 7;276(36):33689–96.

[46] Hong M, Fitzgerald MX, Harper S, et al. Structural basis for dimerization in DNA recognition by Gal4. Structure. 2008 Jul;16(7):1019–26.

[47] Efron B, Hastie T, Johnstone I et al. Least angle regression. Ann. Statist. 2004;32:407–499.

[48] Pedregosa F, Varoquaux G, Gramfort A et al. Scikit-learn: machine learning in Python. J Mach Learn Res 2011;12:2825–30.

[49] Virtanen P, Gommers R, Oliphant TE et al. SciPy 1.0: fundamental algorithms for scientific computing in Python. Nat Methods. 2020;17:261–272.

[50] Rao R, Meier J, Sercu T et al., Transformer protein language models are unsupervised structure learners. International Conference on Learning Representations 2021.

[51] Riesselman, A.J., Ingraham, J.B. & Marks, D.S. Deep generative models of genetic variation capture the effects of mutations. Nat Methods. 2018;15:816–822.

[52] Biswas S, Khimulya G, Alley EC et al. Low-N protein engineering with data-efficient deep learning. bioRxiv doi: 10.1101/2020.01.23.917682.

[53] Barilá D, Superti-Furga G. An intramolecular SH3-domain interaction regulates c-Abl activity. Nat Genet. 1998;18:280–2.

[54] Bradbury J, Frostig R, Hawkins P et al., JAX: composable transformations of Python+NumPy programs. http://github.com/google/jax

[55] Eric J. Ma, Arkadij Kummer. Reimplementing Unirep in JAX. bioRxiv doi: https://doi.org/10.1101/2020.05.11.088344.

[56] Kingma DP, Welling M. Auto-Encoding Variational Bayes. arXiv e-prints, page arXiv:1312.6114, May 2014

